# Nicotinamide riboside: A promising therapy for MI-induced acute kidney injury by upregulating nicotinamide phosphoribosyltransferase-mediated NAD levels

**DOI:** 10.1101/2024.09.05.611567

**Authors:** Nada J. Habeichi, Ghadir Amin, Solene Boitard, Cynthia Tannous, Rana Ghali, Iman Momken, Reine Diab, George W. Booz, Mathias Mericskay, Fouad A. Zouein

## Abstract

**Background:** Cardiorenal syndrome (CRS) type 1 is characterized by the development of acute kidney injury (AKI) following acute cardiac illness and notably acute myocardial infarction (MI). AKI is considered an independent risk factor increasing mortality rate substantially. Nicotinamide dinucleotide (NAD) is an important coenzyme in energy metabolism and oxidative phosphorylation and in its oxidized form, a substrate for multiple NAD^+^-dependent enzymes such as Sirtuins and poly-ADP ribose polymerases. Decreased cardiac NAD levels along with a down-regulation of the nicotinamide phosphoribosyl transferase (NAMPT) have been reported following MI. A compensatory upregulation in nicotinamide riboside kinase (NMRK) 2, an NAD^+^ biosynthetic enzyme that uses nicotinamide riboside (NR) to generate NAD^+^ takes place in the heart after MI but the impact on kidney NAD metabolism and function has not been addressed before.

**Methods:** MI was induced by ligating the left anterior descending coronary artery in 2 months old C57BL6/J mice, followed by the administration of NR (IP injection, 400mg/kg/day) for four and seven days. We hypothesized that NR treatment could be a potential promising therapy for MI-induced AKI.

**Results:** Our findings showed no significant improvement in cardiac ejection fraction following NR treatment at days 4 and 7 post-MI, whereas kidney functions were enhanced and morphological alterations and cell death decreased. The observed renal protection seems to be mediated by an up-regulation of NAMPT-mediated increase in renal NAD levels, notably in distal tubules.

**Conclusion:** Our findings indicate that NR could be a potential promising therapy for AKI following an early stage of MI.

## Introduction

Overwhelming epidemiological evidence links cardiovascular diseases (CVDs) to acute kidney injury (AKI).^1^ According to the World Health Organization (WHO), ischemic heart diseases, including myocardial infarction (MI) and stroke, are responsible for 31% of CVD deaths annually (https://www.who.int/news-room/fact-sheets/detail/cardiovascular-diseases-(cvds). MI is a major public health concern and a leading cause of CVD-induced AKI through the cardiorenal interrelationship, known as type I cardiorenal syndrome (CRS).^2^ Clinical studies demonstrated that between 7 to 29 % of hospitalized MI patients developed AKI and 2.31% will undergo hemodialysis during hospitalization.^3,4,5^ Furthermore, it has been reported that MI patients are not only at a 30-fold increased risk of heart failure progression and development but also have up to a 29-fold increased risk of CRS, which increases short- and long-term mortality rates.^6,7^

CVDs, mainly ischemic heart diseases are associated with a perturbation in cardiac energy metabolism characterized by a shift in energy production from oxidative phosphorylation (OXPHOS) from fatty acid beta-oxidation or glucose oxidation to anaerobic glycolysis when oxygen supply is limited.^8,9^ Nicotinamide adenine dinucleotide (NAD) is emerging as a metabolic target in different CVDs as a major coenzyme in fuel oxidation and OXPHOS, and as a signaling molecule in stress responses through its consumption as a substrate by Sirtuin deacetylases (SIRT-1 to 7) that regulate energy metabolism at different levels and poly(ADPribose) polymerases, notably PARP-1 involved the repair of reactive oxygen species (ROS)-induced DNA damages.^10,11^ Altered NAD levels have been reported in several models of heart diseases including ischemia-reperfusion injury. For instance, a study performed by Di Lisa et al. showed that 30 min of cardiac ischemia resulted in a marked decrease in mitochondrial and tissue NAD^+^ levels in male Wistar rats.^12^ However, little is known about renal NAD homeostasis in the context of type I CRS post-MI. In that regard, our published work showed a significant decrease in kidney NAD levels in 5 months old C57BL6/J male and female mice at day 7 post-MI.^13^ Of note, decreased kidney NAD levels were exacerbated in MI male mice in the presence of risk factors such as cigarette smoking.^14^ Additionally, a marked downregulation in kidney SIRT-1 and SIRT-3 mRNA expression levels along with a substantial upregulation in PARP-1 mRNA expression levels were observed in male mice, only, at day 7 post-MI, suggesting the urgent need for metabolic drugs that can boost NAD levels and correct the observed alteration in NAD signaling pathways.^13^

To date, there are no metabolic drugs that can target type I CRS or limit its development or progression. Nicotinamide riboside (NR), the nucleoside form of vitamin B3, has recently come into focus as an effective NAD precursor with superior bioavailability.^15^ A study performed by Diguet et al. reported a significant decrease in cardiac NAD pools (i.e. the sum of NAD^+^ and NADH) in a mouse model of dilated cardiomyopathy.^16^ NR supplementation (400 mg/kg of body weight/day) attenuated the development and progression of heart failure by rescuing myocardial NAD levels.^16,17^ To date, the potential impact of NR on AKI in the setting of MI is not known. Here we provide for the first time, experimental evidence on the renoprotective impact of NR on AKI at early stage post-MI.

## Methods

### Detailed methods may be found in the Supplementary Material

#### Experimental design

Two months old C57BL6/J male mice were used in this study according to the experimental protocol approved by the Institutional Animal Care and Use Committee (IACUC) of the American University of Beirut (AUB) and in compliance with the National Institutes of Health Guide for the Care and Use of Laboratory Animals, 8th edition.^18^ The approved IACUC protocol number was #18-2-RN560. Mice were randomly divided into 2 groups: SHAM and MI. MI was induced in the morning by ligation of the left anterior descending (LAD) coronary artery and the mice were sacrificed in the morning 4 and 7 days later. The left kidney was immediately placed into a cryotube in liquid nitrogen followed by storage at −80°C for molecular analysis. The right kidney was placed in 4% zinc formalin tubes for histological analysis.

#### Myocardial infarction (MI)

The MI model was established by permanent ligation of the LAD coronary artery. Instant blanching of the LV area below the ligation site and the ST-segment elevation on the echocardiogram indicated successful ligation. The same procedure was performed for SHAM-operated mice but without LAD artery ligation. The mouse was then weaned from isoflurane anesthesia, flushed with oxygen, extubated, and allowed to warm under an infrared lamp until full consciousness.

#### NR administration

Phosphate buffered solution (PBS) vehicle or NR diluted in PBS solution was administrated at a dose of 400 mg/kg of body weight/day in a single intraperitoneal (IP) injection. IP injection was administrated to mice 30 min after the surgery as an immediate intervention and repeated every 24 hours until the day of sacrifice.

#### Echocardiography

Transthoracic echocardiography was performed using the Visual Sonics Echo system (Vevo 2100, VisualSonics, Inc., Toronto, Canada) equipped with a 22–55 MHz (MS550D) linear transducer according to the American Society of Echocardiography guidelines to assess cardiac function. At least three consecutive cardiac cycles were measured and averaged.

### Histological Analysis

#### Periodic acid Schiff (PAS)

The PAS stain was applied on 4 µm thick paraffin-embedded kidney sections to assess changes in the Glomerular corpuscle area and glomerular tuft area, Bowman’s capsule, and proximal convoluted tubule. Slides were mounted and observed under the light microscope at 40× magnification. Measurements were performed using the ImageJ software (https://imagej.nih.gov/ij/).

#### Masson’s Trichrome (MTC)

Masson’s trichrome (MTC) staining was used to determine renal fibrosis. Tissues were observed under the light microscope at 10× magnification and renal fibrosis was measured using ImageJ software.

#### Immunohistochemistry (IHC)

IHC was performed to detect NAMPT protein expression levels in kidney tissues using EnVision FLEX Mini Kit High pH: K8023 (Dako). Slides were scanned on the digital scanner NanoZoomer 2.0-RS (Hamamatsu).

#### TUNEL Assay

FragEL DNA Fragmentation Detection Kit, Colorimetric - TdT Enzyme (Sigma-Aldrish, QIA33) was used to assess DNA fragmentation in kidney tissues according to the manufacturer instructions and slides were analyzed using ImageJ software.

##### RNA Extraction and RT-qPCR

Total RNA was isolated from kidney tissues using TriReagent (Thermo Fisher Scientific, Grand Island, NY, USA). Purity (A260/A280 ratio ≥ 1.8) and concentration were measured using the NanoDrop® ND-1000 UV-Vis Spectrophotometer. Results are expressed as fold changes relative to the mean of individual values obtained in the control SHAM+ vehicle group which was set to 1. Primer sequences used are listed in Table S2.

##### Enzyme-linked immunosorbent assay (ELISA)

Urinary cystatin-C and kidney injury molecule-1 (KIM-1) were assessed using ELISA kits (abcam, ab#201280 and ab#213477). All samples were analyzed in duplicate.

##### NAD extraction and Quantification

Buffer composed of 75% ethanol and 25% HEPES 10 mM pH7.1 was used to extract metabolites from kidney tissues (20 µl/mg tissue). Extracts were diluted (1:20) in water. A volume of 25 µl per well was used (duplicate). A volume of 100 μL of reaction buffer was then added to the extracts. Kinetics of the reaction were followed on an LB 942 Multimode Reader (OD at 550 nm, every 30 s for 20 min). All NAD samples were assayed in duplicate and compared to a range of standard NAD concentrations using linear regression curve equation method between NAD standard concentrations and the slope of the reaction.

##### Statistical analysis

Statistical analysis was performed using GraphPad Prism 7 software. Results are expressed as fold change or mean ± standard error of the mean (SEM). Statistical comparisons were performed using two-way ANOVA followed by Tukey multiple comparisons test. The significance threshold was set at P < 0.05.

## Results

### NR does not preserve cardiac function following acute MI

To validate our model of MI and assess the impact of NR on cardiac dysfunction, echocardiography was performed on mice at day 4 and 7 post-MI. Our data show a comparable decrease in ejection fraction (EF) and cardiac output (CO) at day 4 and 7 post-MI in the presence and absence of NR treatment compared to the SHAM+V groups (Figs. 1A-A′ and B-B′). An interaction (P< 0.05) was detected between NR treatment and cardiac output (CO) only at day 4 suggesting a possible transient protective effect, although this parameter remained lower in MINR group compared to SHAMV group at day 4 and 7. LVESD significantly increased at day 4 and 7 post-MI in the presence and absence of NR treatment when compared to the SHAM+V groups (Figs. S1A-A′). LVEDD significantly increased at day 4 post-MI in the presence and absence of NR treatment when compared to the SHAM+V group (Figs. S1B). A marked increase in LVEDD at 7 days post-MI was observed in the MI+NR group compared to the SHAM+NR group (Figs. S1B′). Of note, the statistical analysis of LVESV, LVEDV, SV, FS, and heart rate is presented in Table S1. In summary, NR given intraperitoneally every 24 h had no protective effect on cardiac function at these early stages post-MI.

**Figure 1:**
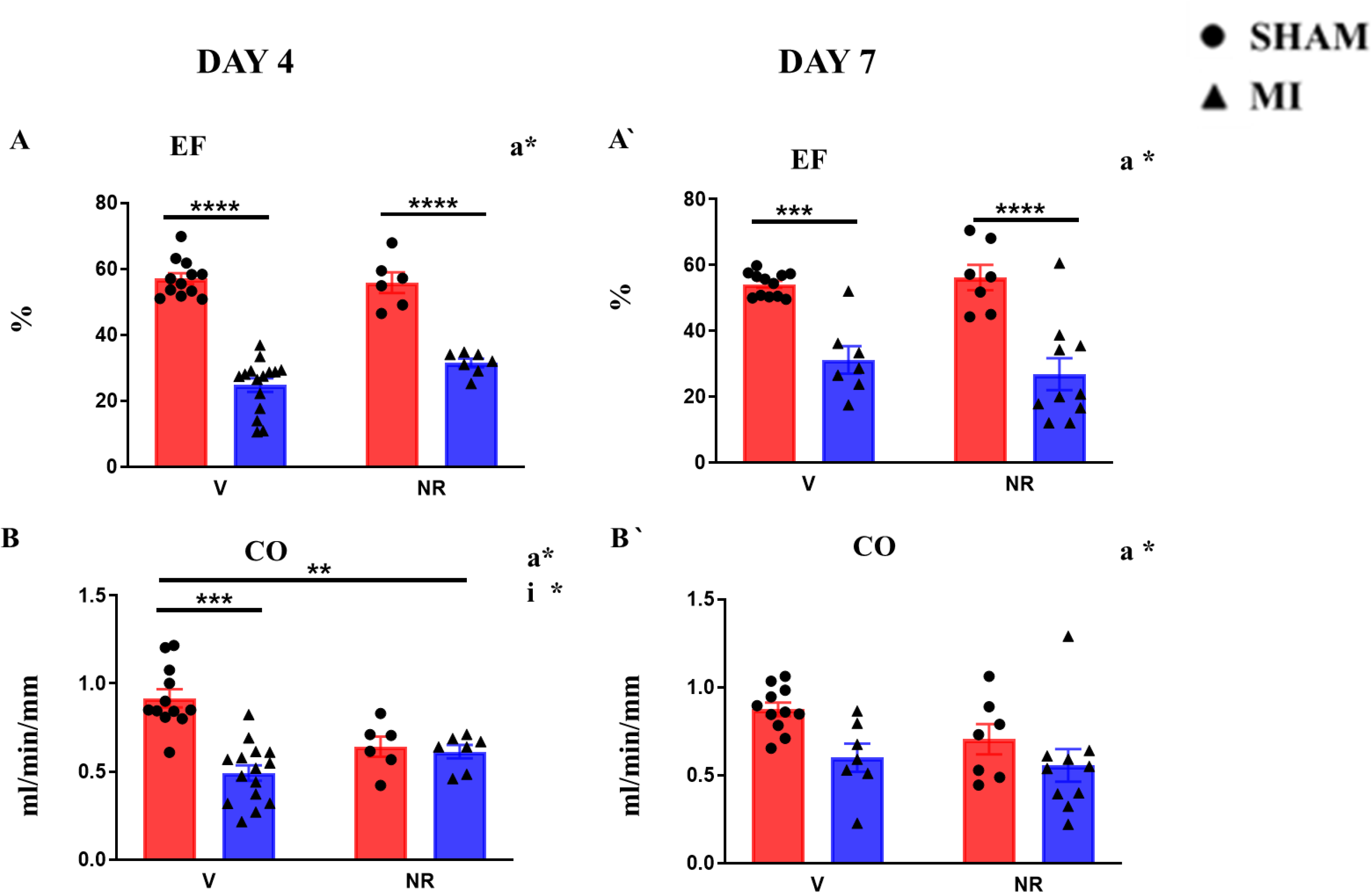
Cardiac function at 4 and 7 days post myocardial infarction with and without NR treatment. A-B: day 4 post-MI, A′-B′: day 7 post-MI. MI: myocardial infarction, V: Vehicle; NR: Nicotinamide riboside. Values are represented as mean ± SEM. Two-Way-ANOVA statistical analysis; a*: MI factor effect <0.05, b*: NR factor effect <0.05, i*: Interaction effect <0.05, n=6-15/group; Tukey post-hoc test: *P<0.05, ***P<0.001, ****P<0.001.

### NR preserves kidney function

To assess the impact of NR on kidney function, we measured urine output, which is recognized as an indicator of exacerbated kidney injury and a predictor of in-hospital mortality.^19^ To further confirm our finding, urinary cystatin-C and kidney injury molecule-1 (KIM-1) were quantified. Cystatin-C and KIM-1 are recognized as diagnostic indicators of enhanced kidney injury and are considered early makers in assessing AKI.^20,21^ MI significantly decreased urine output at day 4 and 7, while it remained unchanged in the MI+NR group (Figs. 2A and A′). Urinary cystatin–C markedly increased at day 4 post-MI, while it remained unchanged following NR treatment. No significant change in urinary cystatin C level was observed at day 7 among all groups (Fig. 2B and B′). NR treatment lowered urinary KIM-1 level in the MI+NR group compared to MI+V group at day 4 post-MI. No significant change in urinary KIM-1 level was observed at day 7 among all groups (Fig. 2 C′). Collectively, our data indicate decreased injury level and improved kidney function at day 4 and 7 post-MI following NR administration compared to MI + V group.

**Figure 2:**
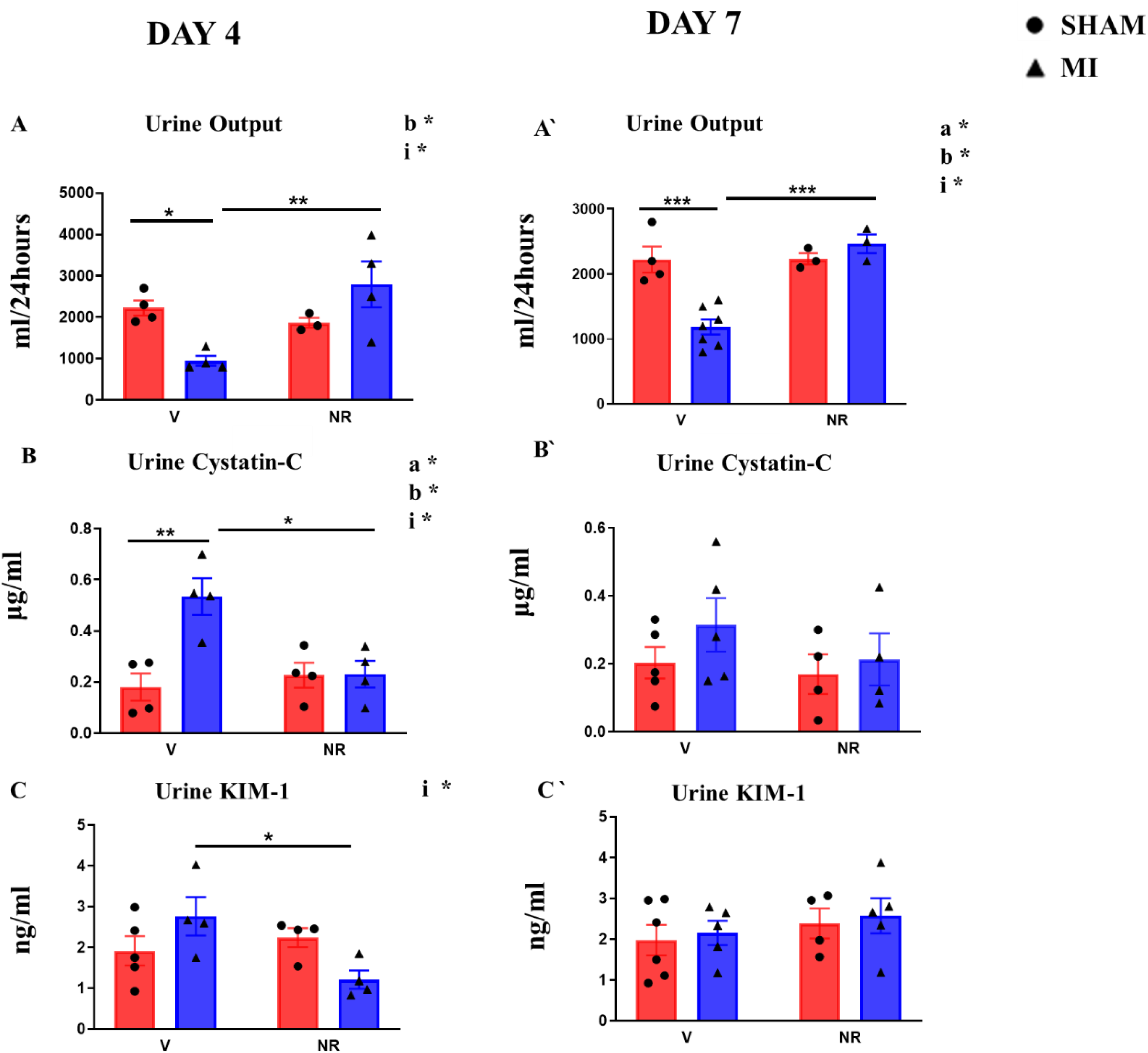
Kidney function at 4 and 7 days post myocardial infarction with and without NR treatment. A-C: day 4 post-MI, A′-C′: day 7 post-MI. KIM-1, kidney injury molecule-1; MI: myocardial infarction, V, vehicle; NR, nicotinamide riboside. Values are represented as mean ± SEM. Two-Way-ANOVA statistical analysis; a*: MI factor effect <0.05, b*: NR factor effect <0.05, i*: Interaction effect <0.05, n=3-7/group; Tukey post-hoc test: *P<0.05, ***P<0.001, ****P<0.001.

### NR reduces kidney morphological alterations

Strong evidence has demonstrated a tight link between ischemic heart diseases including MI and enhanced kidney structural alterations that result in exacerbated kidney dysfunction.^22^ To evaluate the impact of NR on kidney damage, periodic acidic Schiff staining was performed (Figures 3A-A′ and S2A-A′). We observed an increase in glomerular corpuscle area, tuft area, and Bowman space at day 4 post-MI (Figure 3B-D), whereas they remained unchanged following NR treatment. Bowman space was still increased at day 7 post-MI, while it was preserved with NR treatment (Figure 3D′). Proximal convoluted tubule cross-sectional area (PCT-CSA) was substantially enlarged at days 4 and 7 post-MI, but dilation did not take place following NR treatment (Figs. S2B and SB′). Our findings reveal decreased post-MI signs of kidney damage following NR treatment.

**Figure 3:**
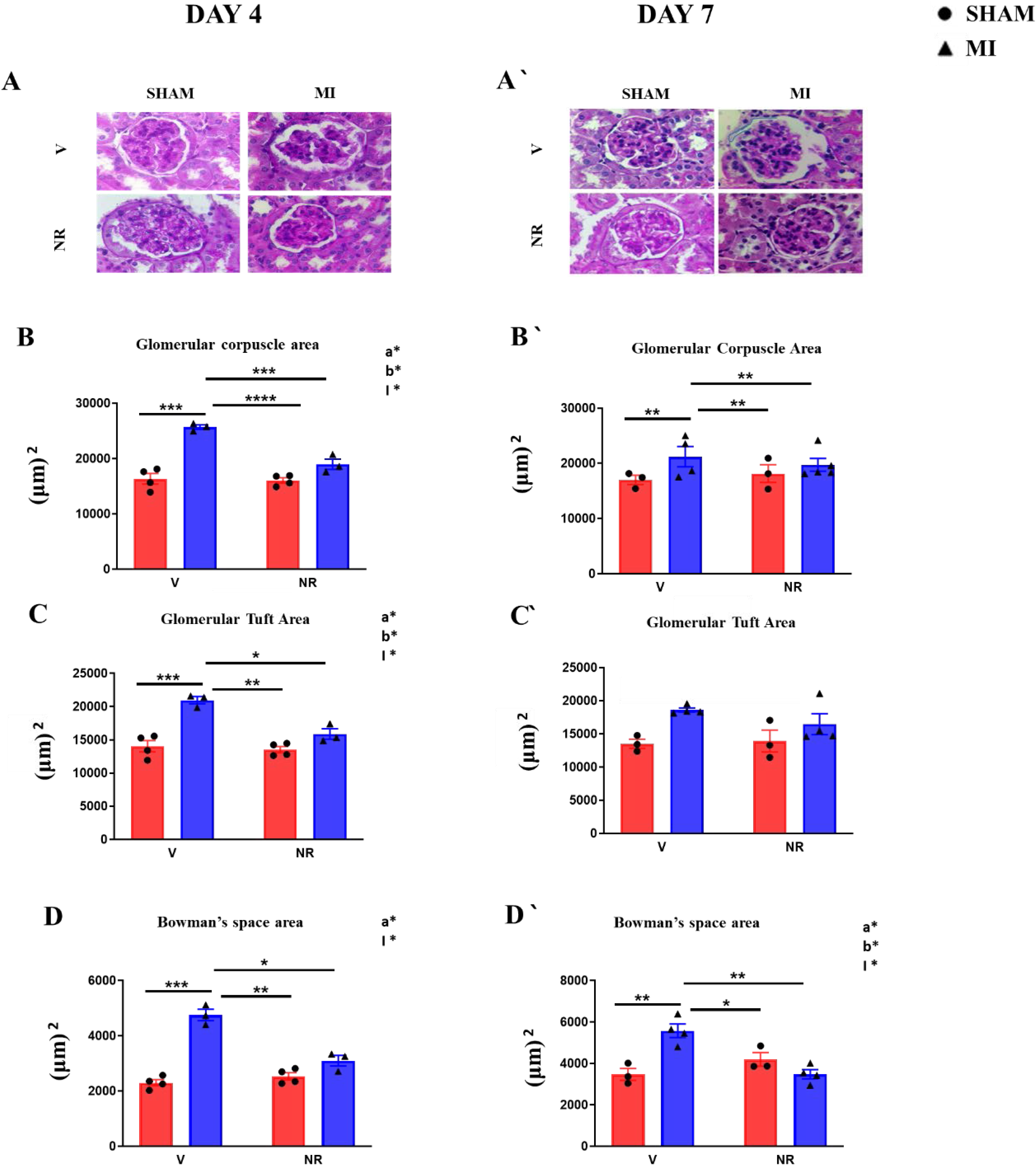
Kidney histology at 4 and 7 days post myocardial infarction with and without NR treatment. A-F: day 4 post-MI, A′-F′: day 7 post-MI. MI, myocardial infarction, V, Vehicle; NR, nicotinamide riboside. Values are represented as mean ± SEM. Two-Way-ANOVA statistical analysis; a*: MI factor effect <0.05, b*: NR factor effect <0.05, i*: Interaction effect <0.05, n=3-5/group; Tukey post-hoc test: *P<0.05, ***P<0.001, ****P<0.001; Scale bar in (A and A′) is 20 µm.

### NR attenuates kidney fibrosis and cell death

AKI is associated with renal fibrosis pathogenesis and enhanced cell death ^23^. In order to assess the impact of NR on kidney fibrosis and cell death, Masson’s trichrome and TUNEL staining were performed (Figure 4A, A′ and C). Our data show a significant increase in renal fibrosis at days 4 and 7 post-MI, whereas it remained unchanged following NR treatment (Fig. 4B and B′). Visualization of TUNEL labeling by IHC showed that cell death particularly increased in the medullary region at day 7 post-MI (Figure 4C). Our findings reveal a protective role of NR on kidney morphological alterations mediated by decreased renal fibrosis and cell death at early-stage post-MI.

**Figure 4:**
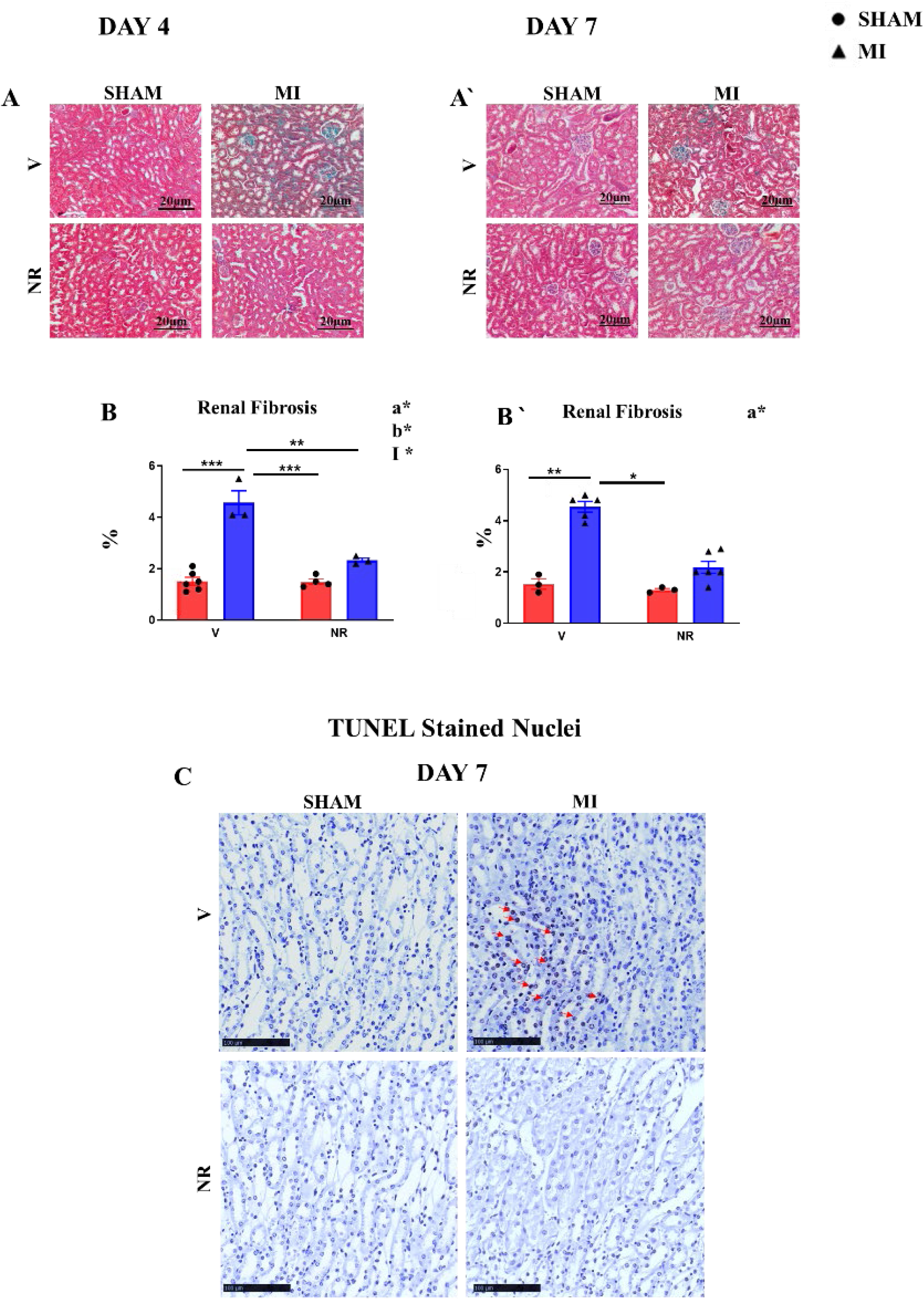
Renal fibrosis and cell death at 4 and 7 days post myocardial infarction with and without NR treatment. A-D: day 4 post-MI, A-D′: day 7 post-MI. A-A′ Masson’s trichrome staining. B-B′: Quantification of fibrosis. MI, myocardial infarction, V, vehicle; NR, nicotinamide riboside. Values are represented as mean ± SEM. Two-Way-ANOVA statistical analysis; a*: MI factor effect <0.05, b*: NR factor effect <0.05, i*: Interaction effect <0.05, n=3-6/group; Tukey post-hoc test: *P<0.05, ***P<0.001, ****P<0.001; Scale bar in (A and A′) is 20 µm and C is 100 µm.

### NR preserves total NAD levels and enhances NAD^+^ signaling pathway

The role of NAD in regulating metabolism and modulating cellular homeostasis is well established.^11,24^ NAMPT and NMRK-1, NAD^+^ biosynthetic enzymes, play a major role in NAD production in the kidneys.^25^ Total NAD levels (sum of NADH and NAD^+^) in the kidneys showed no significant change at 4 and 7 days post-MI in vehicle-treated groups compared to SHAM groups, whereas administration of NR increased its levels in the MI group at both time points but not in the SHAM groups (Figs. 5A and 5A′). No alteration in *Nampt* mRNA expression levels was observed at day 4 and 7 post-MI in vehicle-treated groups, whereas the administration of NR transiently increased its levels at day 4 in the MI group only (Fig. 5B and 5B′). *Nmrk1* mRNA expression showed no significant alteration at days 4 and 7 following MI in the presence and absence of NR treatment (Figs. 5C and 5C′). Our findings indicate that NR administration increased kidney NAD levels as early as 4 and up to 7 days post-MI, and this elevation is associated with an upregulation of *Nampt* mRNA expression levels as soon as day 4.

**Figure 5:**
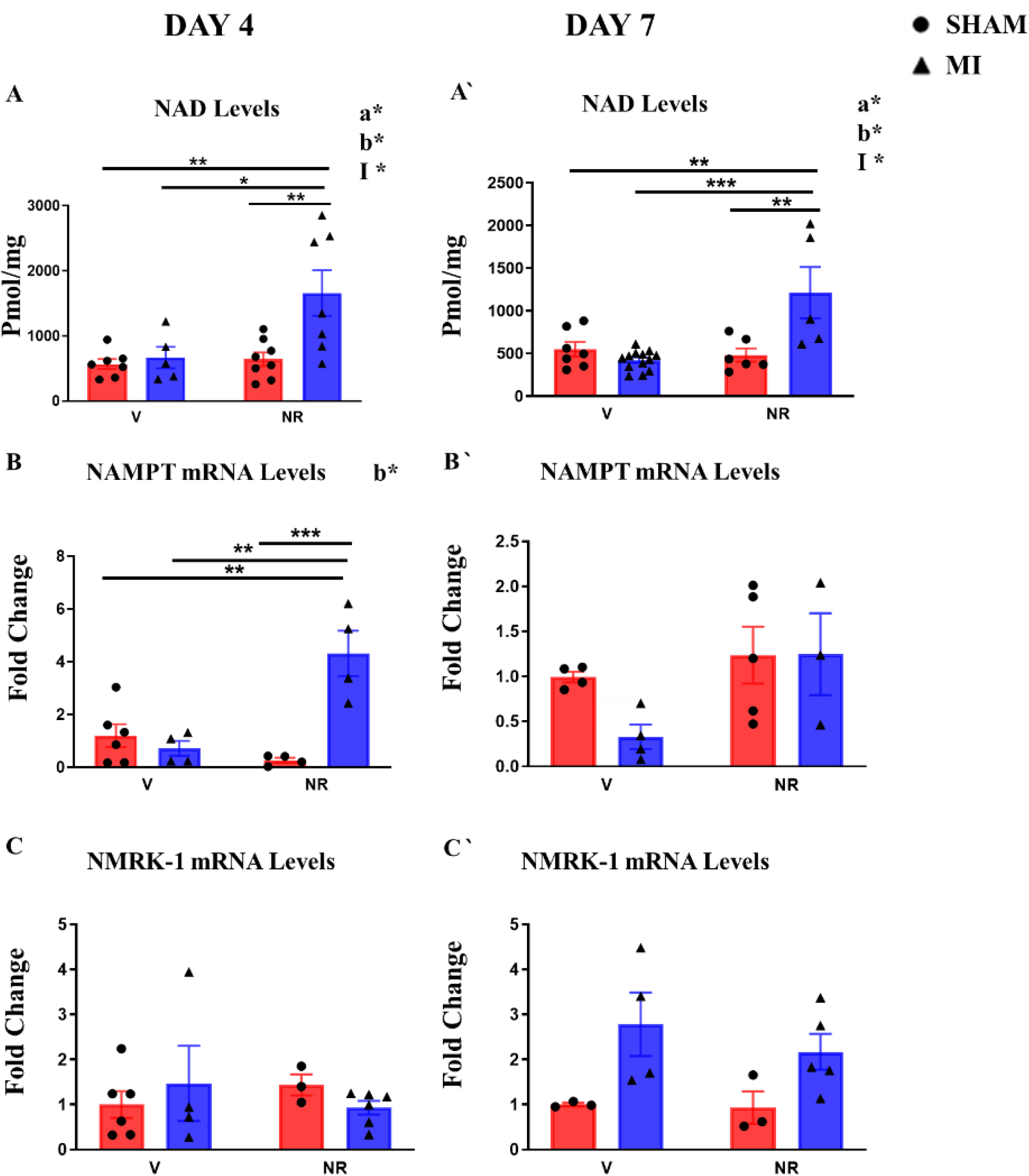
NAD levels and NAD^+^ signaling pathway at 4 and 7 days post myocardial infarction with and without NR treatment. A-C: day 4 post-MI, A′-C′: day 7 post-MI. NAD: nicotinamide adenine dinucleotide, *Nampt*: nicotinamide phosphoribosyltransferase, *Nmrk1*, nicotinamide riboside kinase 1; MI, myocardial infarction, V, vehicle; NR, nicotinamide riboside. Values are represented as mean ± SEM. Two-Way-ANOVA statistical analysis; a*: MI factor effect <0.05, b*: NR factor effect <0.05, i*: Interaction effect <0.05, n=3-13/group; Tukey post-hoc test: *P<0.05, ***P<0.001, ****P<0.001.

Immunohistochemistry was performed to confirm the observed increase in NAMPT expression at day 4 post-MI following NR treatment and to assess in which cell type elevation occurred. In the cortex, NAMPT is present in the nucleus of a few cells located in the glomeruli (Figs. 6A, A′, B and B′, red arrows) and of the vast majority of proximal tubule epithelial cells. In distal tubules, NAMPT is present both in the nucleus and cytosol. Interestingly, in the medullary area which is enriched in distal tubules, NR treatment induced the most striking increase in NAMPT signal (Fig. 6D′).

**Figure 6:**
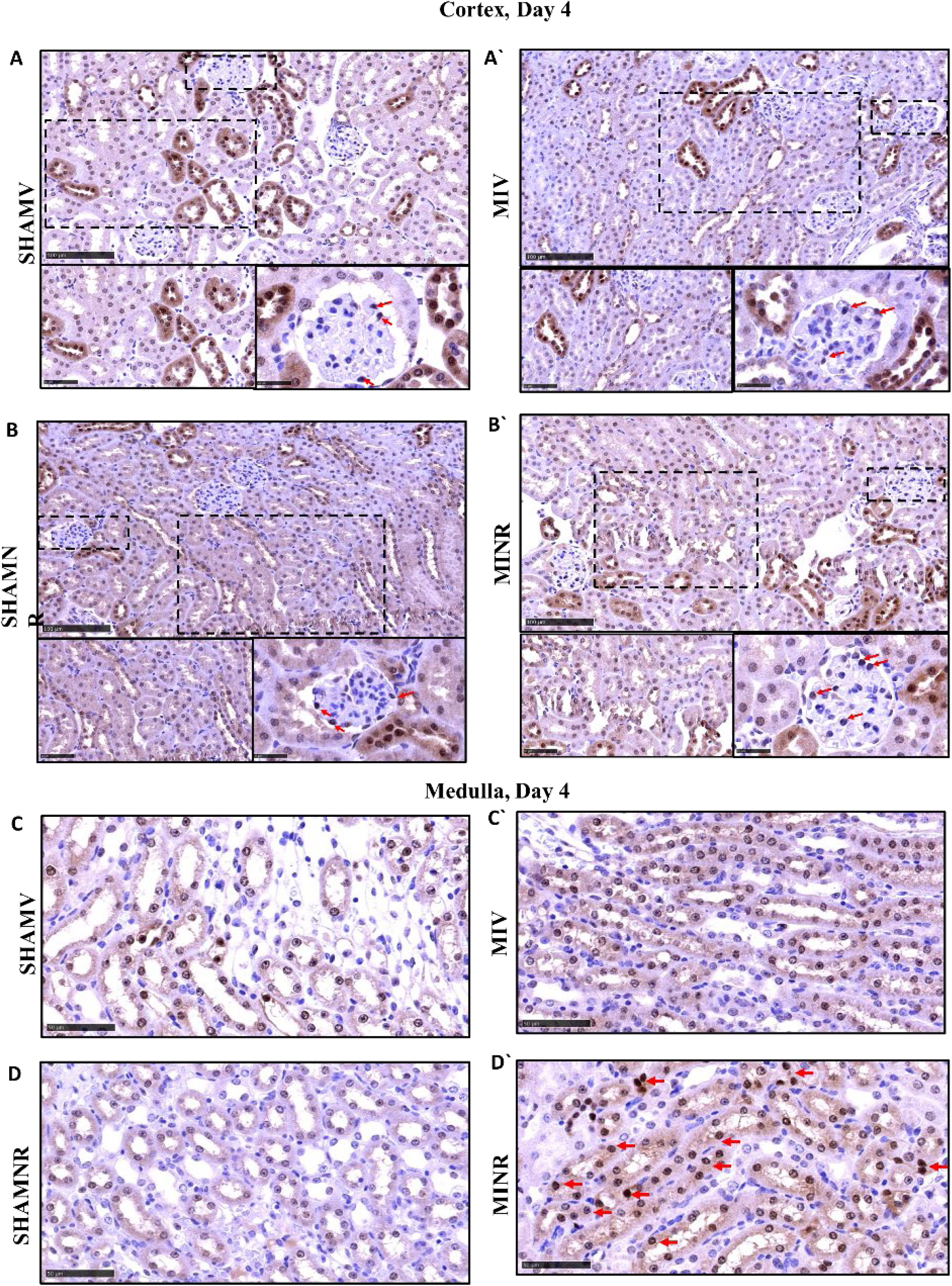
IHC analysis of NAMPT changes in the kidney with NR. Representative images of NAMPT showing an increase in NAMPT protein levels at day 4 following NR treatment at the level of distal tubules. A-A′-B-B′: Cortex region. C-C′-D-D′: Medulla region. n=3/group. Scale bar is 100μm.

## Discussion

Mounting evidence reports a close link between acute cardiac diseases and AKI development and progression.^26–28^ The main objective of this study was to investigate the impact of NR treatment on MI-induced AKI at early-stage post-MI. MI was induced in male mice by ligating the LAD coronary artery followed by sacrifice at either 4- or 7-days post-MI. NR, a NAD precursor, was delivered by intraperitoneal injection (400mg/kg of body weight/day), during a therapeutic window starting 30 min after infarction up to the day of sacrifice (4- and 7-days post-MI).

Strong evidence demonstrates a crucial role of NAD^+^ and NAD^+^ biosynthetic and dependent enzymes in protecting against aggravated oxidative stress and cell death, mitochondrial dysfunction, and enhanced inflammatory response.^29–33^ In this study, we report enhanced kidney dysfunction and morphological alterations and cell death, at early stage post-MI. Post-MI administration of NR, the substrate of NMRK2 in cardiac tissues and NMRK-1 in the kidneys, nearly doubled kidney NAD levels, improved kidney function, decreased kidney morphological alterations, and protected against fibrosis and cell death. As EF and CO were not improved in this MI model following NR treatment for 4 and 7 days, we therefore assume that the decrease in kidney injury could be due to a direct effect of NR on the kidneys.

The alteration of NAD homeostasis has been reported in several experimental models of AKI including ischemia reperfusion injury.^34^ In this study, we observed no change in total NAD pool in the kidneys as early as 4- and 7-days post-MI. This observation could be due to the young age of the mice we chose to use for the present study (2 months old). Our previous work, which had shown a significant decrease in NAD levels in the kidneys at day 7 post-MI was done in 5-month-old mice.^13^ Multiple studies indicate that NAD^+^ metabolism is progressively diminished by aging.^35,36^ Nevertheless, NR administration post-MI still markedly increased NAD levels in the kidneys. Of note, NMRK-1, an enzyme that uses NR as a substrate to generate NAD^+^ tended to increase at day 7 post-MI. On the other hand, NAMPT, an enzyme that uses nicotinamide (NAM) to generate NAD^+^, increased earlier at day 4 post-MI but only following NR treatment. NAMPT protein mainly increased at the level of the medulla. Consistent with this observation, a study performed by Kropotov et al. indicated that NR after entering the cells, is in part metabolized to NAM under the action of purine nucleoside phosphorylase (PNP), leading to the intracellular accumulation of NAM in tissues, including kidneys.^37^ In this study, inhibition of PNP using Immucillin H resulted in increased NAD synthesis from NR and high levels of NR in the kidneys and blood two hours after injection.^37^ To summarize, our data suggest that in the context of AKI following MI, NR can be first metabolized into NAM to generate NAD^+^ through the NAMPT-mediated pathway that is activated by this treatment at day 4 and then probably through a combination of NAMPT and NMRK1 pathways.

In order to reveal if the observed increase in NAD levels following NR administration exerts a protective effect against MI-induced kidney injury, kidney functional, morphological, and molecular parameters were assessed. Our data revealed a decrease in urine output reflecting deterioration in kidney function at day 4 and 7 post-MI. The administration of NR reverses this dysfunction and decreases levels of markers of kidney injury, including KIM-1 and cystatin-C. In that regard, KIM-1 has gained interest as a prognostic maker for renal tubule damage and decline in renal function ^38^. Moreover, urinary cystatin-C has been identified as a better and earlier predictive marker of AKI than creatinine, not being affected by sex, age, or muscle mass.^39,40^

Morphologically, our findings demonstrated aggravated kidney structural alterations at day 4 and 7 post-MI, whereas the administration of NR reversed this damage. In models of kidney injury triggered by ischaemia-reperfusion injury (IRI) or cisplatin injection, administration of NR, 10 days prior to induction of AKI and until the day of sacrifice (400 mg/kg/day diet), was documented to lessen kidney dysfunction and decrease tubular injury analyzed at 48 hours post-renal IRI-induced AKI and 72 hours following cisplatin-induced AKI.^41^ Here, we demonstrate that NR can be administrated as an immediate therapeutic intervention after the MI, establishing a strong potential for translation into clinics in type I CRS. We also found that NR treatment decreased renal fibrosis. In line with this observation, it has been shown that NR administration also attenuates hepatic fibrosis by enhancing the activity of SIRT-1 and supressing expression of P300 ^42^. Furthermore, in a study performed by Pham et al. the administration of NR (400 mg/kg of body weight/day) for 20 weeks decreased liver fibrosis through inhibiting collagen deposition in a mouse model of high fat diet-induced hepatic fibrosis.^43^

## Perspectives

The effects of NR administration in maintaining kidney NAD levels, preserving kidney function and morphology, establish a solid rationale for NR as a promising therapy for lessening AKI pathogenesis in the setting of large MI. Further experiments tailored to evaluate the protein expression levels of NAD biosynthetic and dependent enzymes are warranted. Determining the potential upregulation of PNP in the kidneys following NR administration is of utmost importance. Importantly, in our study we assumed that NR has a direct protective role on the kidneys; however, Freeberg et al. demonstrated that NR could also have an impact on the arteries,^44^ offering a potential additional mechanism for the observed improvement in kidney function in our study. Further investigations, therefore, are needed to reveal the exact molecular pathways that are involved in the observed protection on the kidneys at early stage post-MI.

## Novelty and Relevance

**What is New?**

- NR treatment protects the kidneys during the early stages following MI.
- NR treatment mitigates MI-associated renal structural damage, cellular death, and fibrosis, resulting in improved kidney function.
- Renal protection is facilitated by the upregulation of NAMPT-mediated NAD levels in the distal tubules.

**What is Relevant?**

- Hypertension causes cardiac hypertrophy, a major risk factor for myocardial infarction that may damage kidneys.
- NR protects against MI-associated kidney damage by replenishing the renal NAD pool.

**Clinical Pathophysiological Implications?**

- NR is well tolerated in humans and has no adverse effects on the kidneys or cardiac function in patients with heart failure.
- NR treatment can be considered a potential therapeutic approach to be explored in a controlled clinical trial for type I CRS.

## Acknowledgements

We thank Françoise Mercier-Nomé for valuable training and help with immunohistological analyses in the histopathology core (PHIC) and the support of the Animex team (animal experimentation); both are facilities of the “Ingénierie et Plateformes au Service de l’Innovation Thérapeutique” UMS IPSIT of Université Paris-Saclay - US 31 INSERM - UAR3679 CNRS. GWB acknowledges the support of the Pharmacology Clinical Research Core of the University of Mississippi Medical Center.

## Non-standard Abbreviations and Acronyms

AKI: Acute kidney injury
CO: Cardiac output
CRS: Cardiorenal syndrome
CVDs: Cardiovascular diseases
EF: Ejection fraction
FS: Fractional shortening
LAD: Left anterior descending
IACUC: Institutional Animal Care and Use Committee
IHC: Immunohistochemistry
KIM-1: kidney injury molecule-1
LVEDD: Left ventricular end-diastolic diameter
LVESD: Left ventricular end-systolic diameter
LVEDV: left-ventricular end-diastolic volume
LVESV: Left-ventricular end-systolic volume
MI: Myocardial infarction
MTC: Masson’s trichrome
NAD: Nicotinamide dinucleotide
NAMPT: Nicotinamide phosphoribosyl transferase
NMRK: Nicotinamide riboside kinase
NR: Nicotinamide riboside
OXPHOS: Oxidative phosphorylation
PBS: Phosphate buffered solution
ROS: Reactive oxygen species
SV: Stroke volume

---

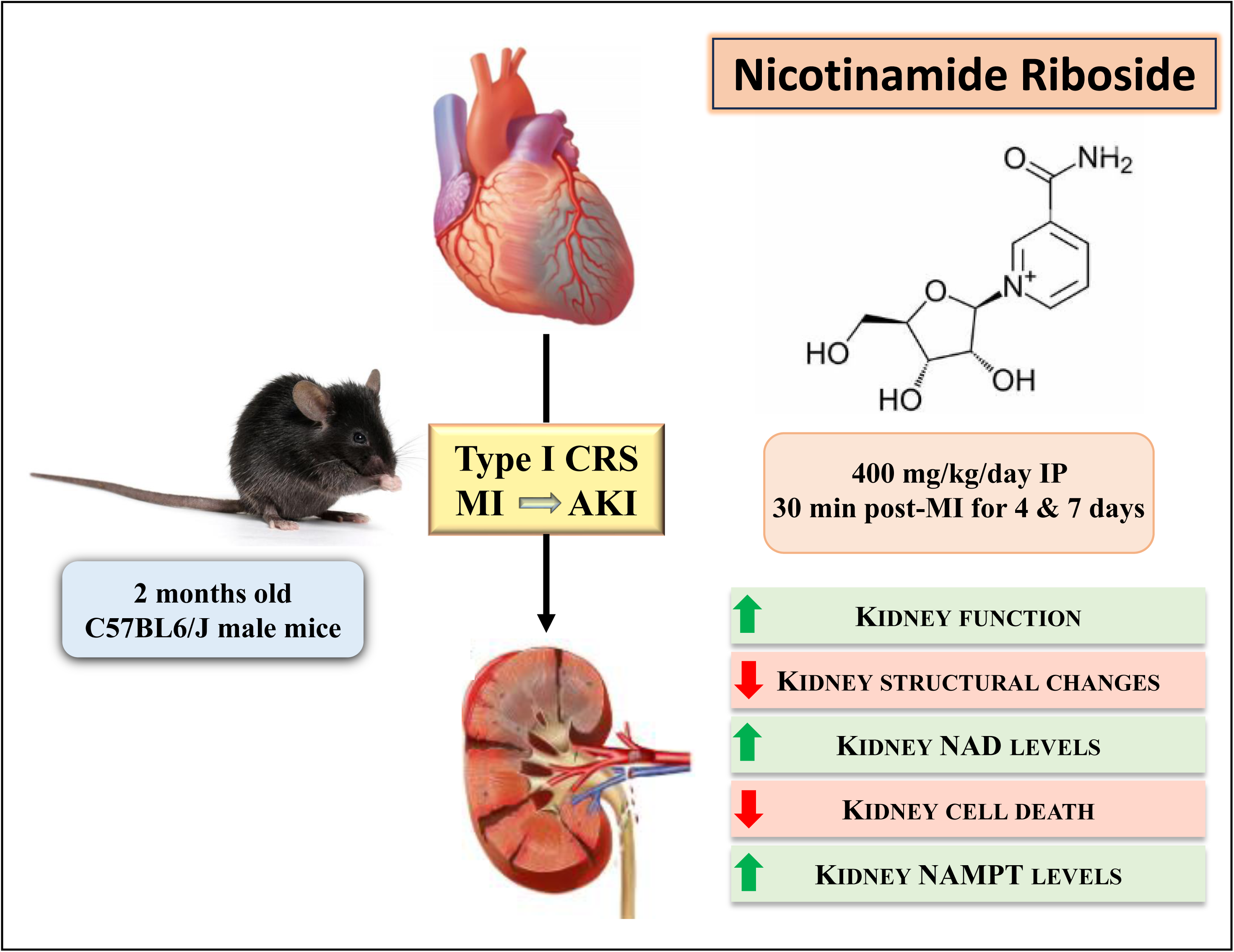

**Figure S1.**
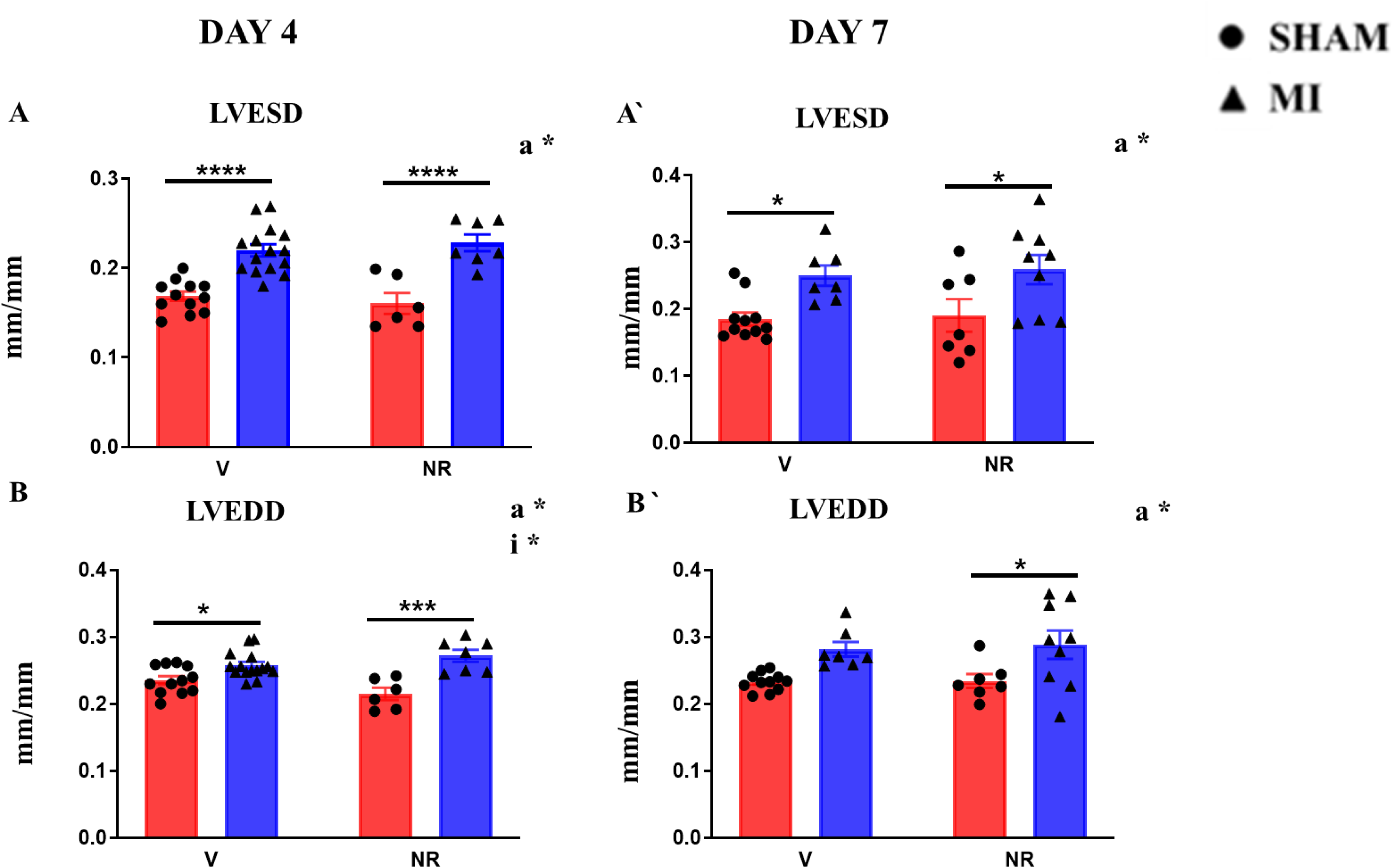

**Figure S2.**
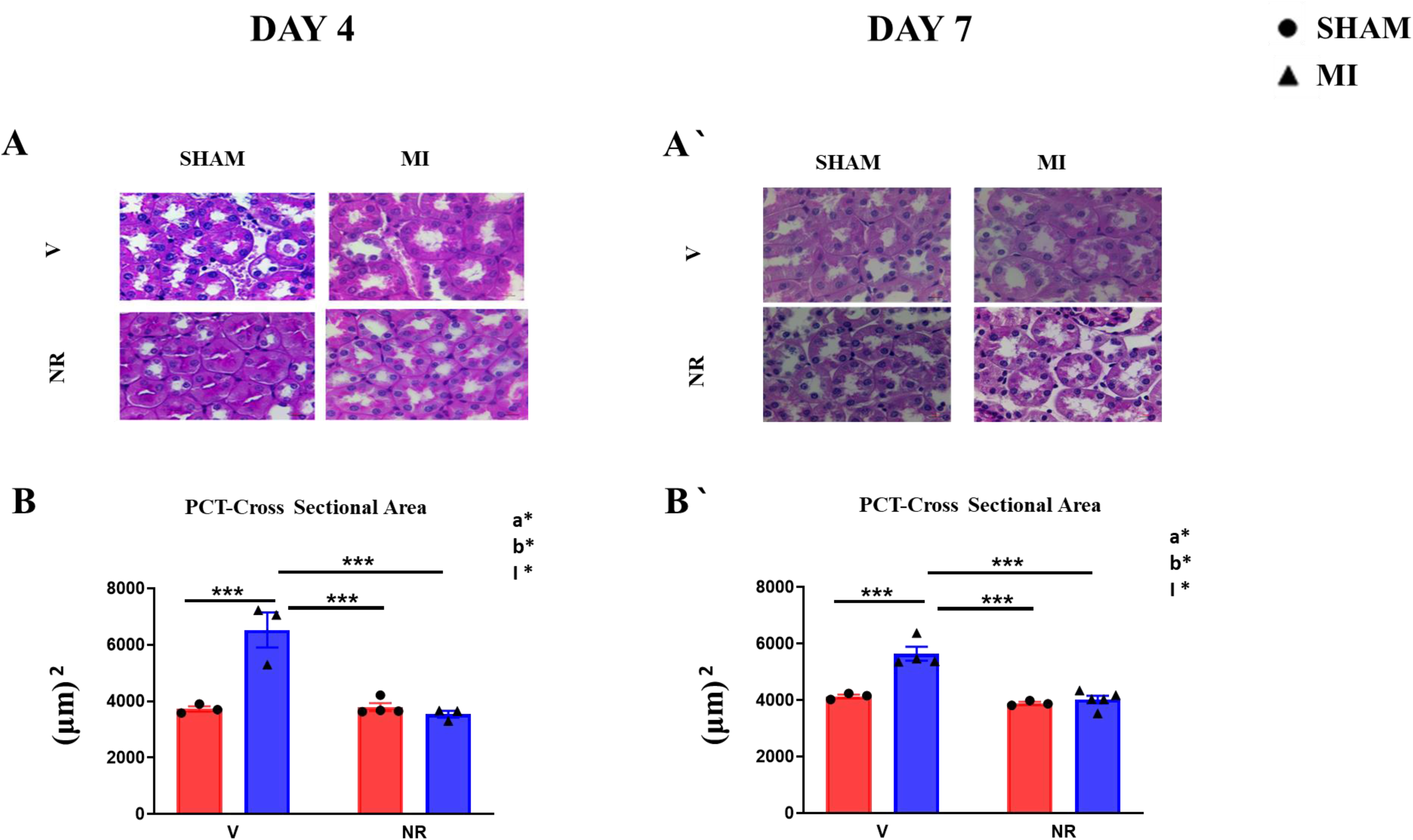

